# The Astrin-SKAP Complex Reduces Friction at the Kinetochore-Microtubule Interface

**DOI:** 10.1101/2021.11.29.469773

**Authors:** Miquel Rosas-Salvans, Renaldo Sutanto, Pooja Suresh, Sophie Dumont

## Abstract

The kinetochore links chromosomes to spindle microtubules to drive chromosome segregation at cell division. While we know nearly all mammalian kinetochore proteins, how these give rise to the strong yet dynamic microtubule attachments required for function remains poorly understood. Here, we focus on the Astrin-SKAP complex, which localizes to bioriented kinetochores and is essential for chromosome segregation, but whose mechanical role is unclear. Live imaging reveals that SKAP depletion dampens movement and decreases coordination of metaphase sister kinetochores, and increases tension between them. Using laser ablation to isolate kinetochores bound to polymerizing vs depolymerizing microtubules, we show that without SKAP kinetochores move slower on both polymerizing and depolymerizing microtubules, and that more force is needed to rescue microtubules to polymerize. Thus, in contrast to previously described kinetochore proteins that increase grip on microtubules under force, Astrin-SKAP reduces grip, increasing attachment dynamics and force responsiveness and reducing friction. Together, our findings suggest a model where the Astrin-SKAP complex effectively “lubricates” correct, bioriented attachments to help preserve them.

## INTRODUCTION

The kinetochore links each chromosome to spindle microtubules at cell division, transmitting spindle forces to move chromosomes. To perform its function, the kinetochore must not only bind microtubules strongly enough to resist cellular forces, but also slide on them to move and segregate chromosomes. While we now have a detailed map of mammalian kinetochore components, and are uncovering their structure, biochemistry and biophysics, how these components together give rise to the mechanics of the kinetochore-microtubule interface remains poorly understood. Indeed, we cannot as yet reconstitute mammalian kinetochores or the microtubule bundles they bind to *in vitro*, and applying precise mechanical perturbations to mammalian kinetochores remains challenging *in vivo*. How mammalian kinetochore-microtubule attachments can be robust and strong yet dynamic remains an open question. Answering this question is central to understanding how cells accurately segregate their chromosomes.

To perform its function, the kinetochore-microtubule interface both generates and responds to force. In mammalian cells, kinetochores bind to the 15-25 microtubules that form a kinetochore-fiber (k-fiber) (Wendell *et al*, 1993; McEwen *et al*, 1997), and that both polymerize and depolymerize. When sister kinetochores oscillate together at metaphase, active (energy consuming) force generation from microtubule depolymerization at the “front” kinetochore (moving towards the pole) largely drives movement of the pair; in turn, passive, frictional force at the “back” kinetochore (moving away from the pole) is generated as kinetochore proteins slide on microtubules and oppose movement (Figure 1A) (Khodjakov & Rieder, 1996; Grishchuk & McIntosh, 2006; Joglekar *et al*, 2010; Dumont *et al*, 2012; Wan *et al*, 2012; Rago & Cheeseman, 2013). The kinetochore-microtubule interface also responds to force. For example force coordinates microtubule dynamics at both sister kinetochores as chromosomes move (McNeill & Berns, 1981; Skibbens *et al*, 1995; Wan *et al*, 2012), and helps maintain chromosomes in the spindle center through spatially regulated polar ejection forces (Rieder & Salmon, 1994; Ke *et al*, 2009). Key to the interface’s ability to generate and respond to force, it is dynamic: this allows kinetochore mobility on microtubules and microtubule growth and shrinkage, and as such perturbing microtubule dynamics causes segregation defects (Bakhoum & Compton, 2012). While we now know different kinetochore molecules that generate force and increase grip at the microtubule interface (DeLuca *et al*, 2004; Cheeseman & Desai, 2008; Schmidt *et al*, 2012; Zaytsev *et al*, 2014; Long *et al*, 2017; Auckland *et al*, 2017; Huis In ’T Veld *et al*, 2019; Long *et al*, 2019), the mechanisms that make this interface dynamic and able to respond to force despite this grip remain poorly defined.

**Figure 1.**
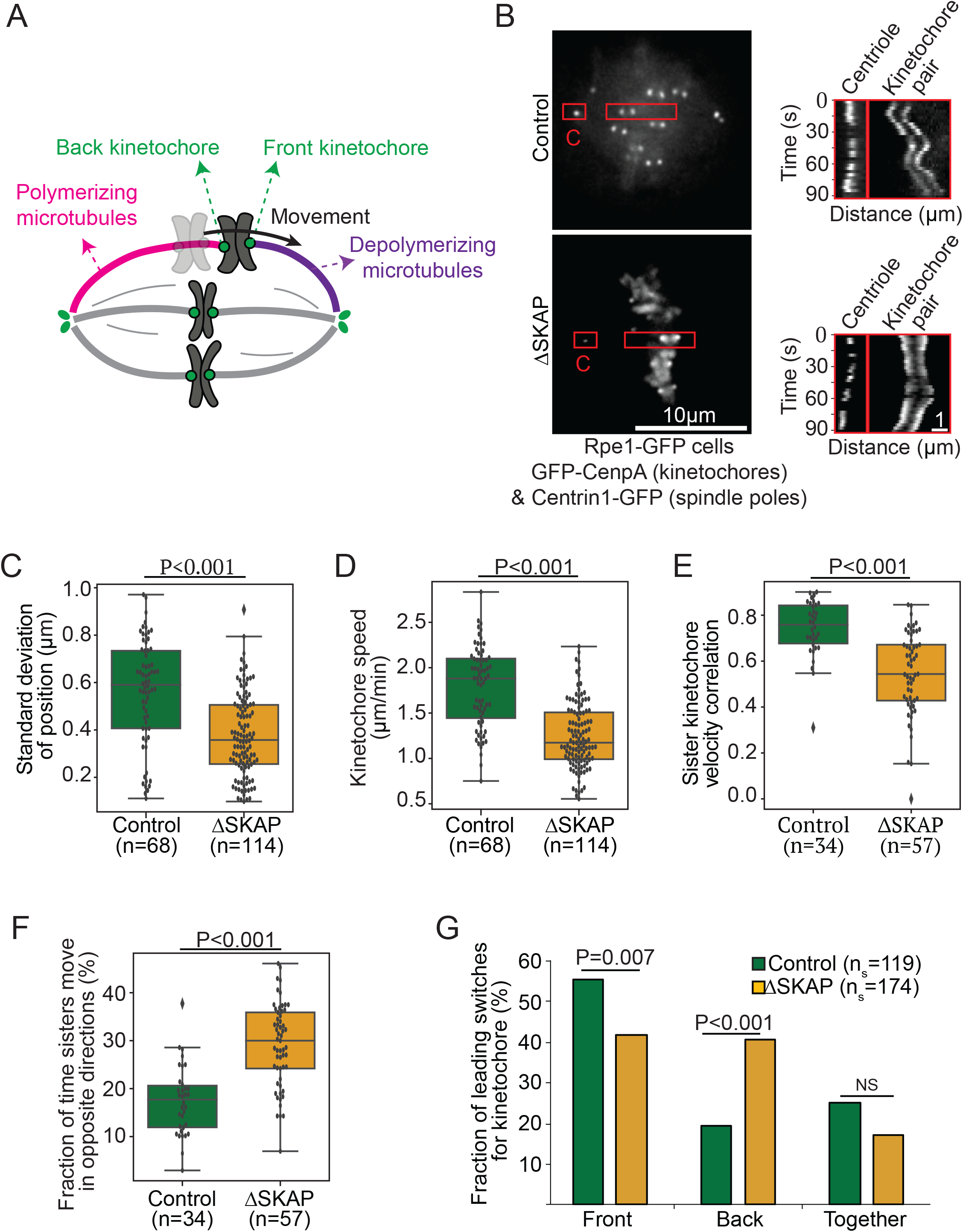
SKAP increases kinetochore mobility and is essential for sister kinetochore coordination in metaphase. (A) Simplified representation of metaphase chromosome oscillations. Force from depolymerizing microtubules (purple) at the front kinetochore drives movement of both sisters. Frictional force at the back kinetochore, bound to polymerizing microtubules (pink), opposes movement. (B) Representative live images (left) of a control and ΔSKAP metaphase Rpe1-GFP cell (GFP-CenpA and centrin1-GFP) with red boxes highlighting regions used for kymographs (right) of centriole (C) and kinetochore movements for those cells. (C) Standard deviation of the position of individual control and ΔSKAP metaphase kinetochores over time (Mann-Whitney test). (D) Average speed of individual control and ΔSKAP metaphase kinetochores (Mann-Whitney test). (E) Velocity correlation between metaphase sister kinetochores (Mann-Whitney test). (F) Fraction of time individual metaphase sister kinetochores move in opposite directions (Mann-Whitney test). (G) Fraction of metaphase directional switches in which the front or back kinetochore switches first, or both switch together (Fisher’s exact test) (n_s_=number of switches). (C-G) from individual kinetochore tracks obtained from the dataset as (B) (n=number of kinetochore pairs, 1-4 kinetochore pairs per analyzed cell from 18 control and 20 ΔSKAP cells). See also Figure S1, Movies S1-2.

Ndc80 and Ska kinetochore complexes play central roles in microtubule attachment in mammalian cells. The Ndc80 complex is essential to the formation of kinetochore-microtubule attachments *in vivo* (DeLuca *et al*, 2004) and directly binds microtubules forming load-bearing attachments *in vitro* (Cheeseman *et al*, 2006; Powers *et al*, 2009). During mitosis, progressive dephosphorylation of the Ndc80 component Hec1 increases its microtubule affinity (Zaytsev *et al*, 2015), load-bearing ability (Huis In ’T Veld *et al*, 2019), and grip on polymerizing microtubules (Zaytsev *et al*, 2014; Long *et al*, 2017). Thus, Ndc80 binds more stably to microtubules as attachments mature, with one phosphorylation site (S69 on protein Hec1) maintaining basal interface dynamics (DeLuca *et al*, 2018). In turn, the Ska complex is essential for proper chromosome alignment and mitotic progression, but is only loaded at kinetochores once these biorient, and in a Ndc80-dependent manner (Hanisch *et al*, 2006). *In vivo*, Ska increases attachment stability to depolymerizing microtubules under force (Auckland *et al*, 2017). *In vitro*, Ska directly binds microtubules (Welburn *et al*, 2009), and increases Ndc80’s affinity for microtubules, and microtubule tracking and load-bearing ability on depolymerizing microtubules (Schmidt *et al*, 2012; Helgeson *et al*, 2018; Huis In ’T Veld *et al*, 2019). Thus, Ska is thought to be a “locking” factor increasing grip on microtubules and stabilizing mature attachments. In addition to Ndc80 and Ska, the Astrin-SKAP (SKAP for short thereafter) complex has been proposed to contribute to microtubule attachment. Like Ska, SKAP is essential for chromosome alignment and mitotic progression, and is only loaded at bioriented kinetochores in a Ndc80-dependent manner (Fang *et al*, 2009; Schmidt *et al*, 2010; Dunsch *et al*, 2011). Strikingly, similar to Ndc80 dephosphorylation (Cimini *et al*, 2006; Long *et al*, 2017), SKAP depletion decreases k-fiber poleward flux (Wang *et al*, 2012), suggesting that SKAP’s presence at the kinetochore may not increase grip on microtubules. Yet, SKAP directly interacts with microtubules *in vitro* (Friese *et al*, 2016; Kern *et al*, 2017), synergistically with Ndc80 (Kern *et al*, 2017). Similar to Ndc80 and Ska (Gao *et al*, 2021; Yu *et al*, 2021), mutations or changes in SKAP expression are highly frequent in some cancers and increase aneuploidy (Siprashvili *et al*, 2014; Deng *et al*, 2021). Yet, SKAP’s mechanical role at the kinetochore-microtubule interface, if any, is not known. More broadly, whether all microtubule-binding kinetochore proteins increase microtubule grip, as Ndc80 and Ska complexes do, or whether the presence of some proteins instead reduce grip to “lubricate” the attachment, remains an open question.

Here, we show that SKAP decreases grip at the kinetochore-microtubule interface, effectively “lubricating” it. We use live imaging to show that SKAP increases the magnitude of metaphase kinetochore movements and the coordination between sisters, and yet that it decreases the tension sisters are under. Using laser ablation, we show that SKAP increases the velocity with which kinetochores move on both polymerizing and depolymerizing microtubules, and that it makes the dynamics of attached microtubules more force-responsive. Thus, not all kinetochore proteins increase the grip on microtubules: SKAP does exactly the opposite, reducing friction at the interface. We propose that SKAP promotes accurate segregation by “lubricating” correct, mature attachments, ensuring that they can smoothly slide despite mechanisms that stabilize them late in mitosis. More broadly, our work suggests that maintaining a strong yet dynamic kinetochore-microtubule interface not only requires components that grip – which are being actively studied – but components that help slide, distinct from those that grip.

## RESULTS

### SKAP increases kinetochore mobility and is essential for sister kinetochore coordination in metaphase

To probe the mechanical role of the Astrin-SKAP complex at the kinetochore-microtubule interface, we live imaged metaphase chromosome movements in human Rpe1 GFP-CenpA (kinetochore) and centrin1-GFP (centriole) cells (Rpe1-GFP cells thereafter (Paul *et al*, 2011)). Upon SKAP depletion by RNAi (Δ SKAP, Figure S1A-B), kinetochore pairs oscillated less far about their mean position than control (0.4±0.2 μm vs 0.6±0.2 μm standard deviation, Figure 1B-C, Movie S1-2), as expected (Wang *et al*, 2012). Δ SKAP kinetochores moved at slower velocity than control (1.2±0.4 μm/min vs 1.8±0.4 μm/min, Figures 1B, D, Movies S1-2), indicating that SKAP increases kinetochore mobility. This suggests that SKAP increases kinetochore-microtubule interface dynamics. Additionally, sister kinetochore movement coordination decreased without SKAP. Δ SKAP sister kinetochore pairs showed lower velocity correlation than control sister pairs (0.54±0.18 vs 0.74±0.12, Figure 1E), with Δ SKAP sister kinetochores moving in opposite directions a higher fraction of time than control (29±7% vs 17±7%, Figure 1F). Further, while in control sister kinetochores the front kinetochore usually reverses direction before the back kinetochore (Figure 1G and Figure S1C (Wan *et al*, 2012)), in Δ SKAP sisters this preference was lost and the back kinetochore switched first more often than control (41% vs 19%, Figure 1G). Together, these findings indicate that SKAP is essential for sister kinetochore mobility and coordination at metaphase.

### SKAP decreases tension at the kinetochore-microtubule interface

Mechanical force from the sister kinetochore and from the spindle on chromosome arms is thought to coordinate sister kinetochore movement and inform directional kinetochore switching (Khodjakov & Rieder, 1996; Ke *et al*, 2009; Wan *et al*, 2012). To probe if miscoordination in Δ SKAP sister kinetochore movement (Figure 1E-G) could be due to defects in force generation or in how the kinetochore respond to force, we measured the interkinetochore (K-K) distance in Δ SKAP cells. Decreasing the activity of microtubule-binding kinetochore proteins (such as Hec1 phosphorylation or Ska depletion) typically reduces the kinetochore’s grip or ability to sustain force before sliding on microtubules, and thus leads to a lower K-K distance (DeLuca *et al*, 2006; Raaijmakers *et al*, 2009; Zaytsev *et al*, 2014) (Figure 2A). In contrast, we found that in live Rpe1-GFP cells Δ SKAP kinetochore pairs had a higher K-K distance than control (1.4±0.2 μm vs 1.1±0.1 μm, Figure 2B), consistent with some (Schmidt *et al*, 2010) but not other (Fang *et al*, 2009; Huang *et al*, 2011) previous reports.

**Figure 2.**
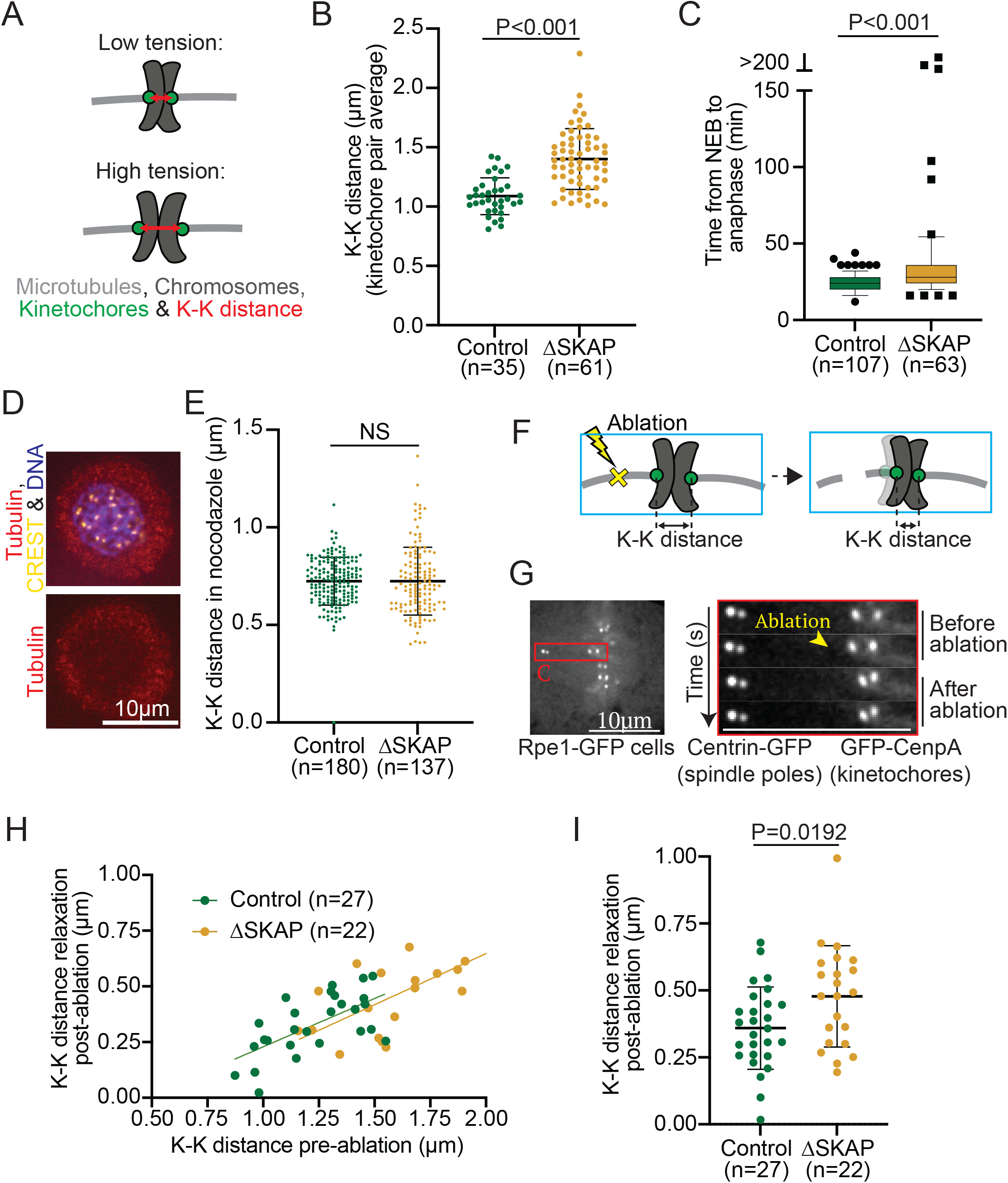
SKAP decreases tension at the kinetochore-microtubule interface. (A) High tension between sister kinetochores leads to a high K-K distance (red double arrow). High tension can stem from high spindles forces, a tighter grip of kinetochores on spindle microtubules, or both. (B) K-K distance average over time for individual kinetochore pairs in control and ΔSKAP Rpe1-GFP cells from the dataset in Figure 1 (student’
ss t-test) (n=number of kinetochore pairs, 1-4 kinetochore pairs per analyzed cell from 18 control and 20 ΔSKAP cells). (C) Time that individual control or ΔSKAP cells spend from nuclear envelope breakdown (NEB) to anaphase onset (Mann-Whitney test) (n=number of cells). (D) Representative immunofluorescence images in control and ΔSKAP nocodazolde treated Rpe1-GFP cells (2µM nocodazole, 3 h) stained for CREST (yellow), chromosomes (purple) and tubulin (red). (E) K-K distance for individual sister pairs in control and ΔSKAP cells treated with nocodazole (Mann-Whitney test) (n=number of kinetochore pairs). (F-G) Laser ablation (yellow X) of k-fiber near a kinetochore releases tension, if present, across a sister pair (schematic cartoon) (F), as shown in representative kymograph images of K-K distance relaxation upon k-fiber ablation (yellow arrowhead) in Rpe1-GFP cells (C for centrioles) (G). (H-I) K-K distance relaxation (decrease) post-ablation as a function of K-K distance pre-ablation (linear regression lines for each condition) (H) or as a direct comparison (I) (student’s t-test) in control vs ΔSKAP cells (n=number of ablations, one ablation per cell). See also Figure S2.

In principle, an increase in K-K distance could stem from increased tension at the kinetochore-microtubule interface, or decreased centromere stiffness (Daum *et al*, 2011; Wan *et al*, 2012). To test whether Δ SKAP cells could arrest at mitosis, and thereby indirectly undergo cohesion fatigue and centromere softening (Daum *et al*, 2011), we imaged and measured mitotic duration (nuclear envelope breakdown to anaphase onset) in Rpe1-GFP cells. Mitotic duration was ∼24 min in control and ∼8 min longer in Δ SKAP cells (23.9±5.8 min vs 31.4±15.0 min, Figure 2C). However, when we artificially induced mitotic arrest using the proteasome inhibitor MG132 (10μm), the K-K distance only detectably increased starting 106 min post-MG132 (Figure S2A-B (Daum *et al*, 2011)). Considering that just 6% of Δ SKAP cells had a mitotic duration longer than 100 min (Figure 2C), cohesion fatigue is unlikely responsible for increasing K-K distance in Δ SKAP cells. To test whether SKAP could more directly affect centromere stiffness, we compared chromosome movements in these MG132-treated cells to Δ SKAP cells (Figure S2). MG132-treated cells selected for their high mean K-K distance (Figure S2A), indicative of low centromere stiffness, had poor sister kinetochore velocity correlation, indistinguishable from Δ SKAP cells, but kinetochore velocity indistinguishable from control (Figure S2C-D). Thus, SKAP cannot simply increase centromere stiffness. Consistent with SKAP not affecting centromere stiffness, treating Rpe1-GFP cells with nocodazole to remove microtubules and spindle forces led to reduced K-K distances that were indistinguishable between Δ SKAP and control (0.7±0.2 μm Δ SKAP vs 0.7±0.1μm control, Figure 2D-E). Thus, the increased K-K distance in Δ SKAP kinetochores depends on microtubules. Supporting this idea, laser ablating k-fibers in Rpe1-GFP cells to release spindle forces (Figure 2F-G) indicates that Δ SKAP kinetochores are under more tension than control. As expected for a spring, sister pairs with higher K-K distances pre-ablation relaxed more post-ablation than those with lower K-K distances in both control and Δ SKAP cells (Figure 2H). Notably, K-K distance relaxed more post-ablation in Δ SKAP kinetochores than control (0.5±0.2 μm vs 0.4±0.1 μm, Figure 2I). This difference indicates that Δ SKAP kinetochores are under higher tension at the kinetochore-microtubule interface, and suggests that they do not slide as much on microtubules under force; this is consistent with sister kinetochores coordinating movement more poorly (Figure 1E-G). Thus, SKAP reduces tension at the kinetochore-microtubule interface, and not all microtubule couplers at the kinetochore increase tension at this interface.

### SKAP decreases kinetochore friction on polymerizing microtubules

To decrease tension at the kinetochore-microtubule interface, SKAP could either reduce passive, frictional force, or active, energy consuming force generated at this interface, or both. At metaphase, the back sister kinetochore is typically bound to polymerizing microtubules through a largely passive interface, and the front sister to depolymerizing microtubules through an interface that is both active and passive (Dumont *et al*, 2012). Thus, decoupling the roles of SKAP at passive versus active interfaces requires uncoupling the effect of SKAP depletion at kinetochores in polymerizing and depolymerizing microtubules.

Since both sisters are attached together, exerting force on each other and holding on to microtubules in opposite polymerization states, we turned to laser ablation to decouple their responses. To probe kinetochores bound to polymerizing microtubules and generating passive force, we ablated k-fibers to trigger spindle-based force generation on a sister pair (Long *et al*, 2017). K-fiber ablation generates new microtubule minus-ends that are recognized by dynein and pulled to the spindle pole (Figure 3A), which exerts an “external” force on the attached sister pair and induces microtubule polymerization at the back (away from the pole) kinetochore (Sikirzhytski *et al*, 2014; Elting *et al*, 2014). Assuming that this dynein pulling force is comparable between conditions, the velocity of this back kinetochore reports on the passive, frictional force at its microtubule interface (Long *et al*, 2017). We ablated k-fibers in Rpe1-GFP cells with and without SKAP (Figure 3B, Movie S3-4), and tracked both sisters. The velocity of the front kinetochore was indistinguishable in Δ SKAP and control cells (3.2±1.5 μm/min Δ SKAP vs 3.0±1.4 μm/min in controls, Figure 3C, E). This is consistent with the dynein-generated force not being affected in Δ SKAP cells. In contrast, the back kinetochore moved slower in Δ SKAP cells than control (2.0±0.9 vs 2.8±1.2 μm/min, Figure 3D-E). While front and back kinetochores moved at indistinguishable velocities in control cells (3.0±1.4 μm/min in front vs 2.8±1.2 μm/min in back kinetochores), back kinetochores moved at lower velocities than front ones in Δ SKAP cells (2.0±0.9 μm/min vs 3.2±1.5 μm/min) (Figure 3E). This led to a persistent increase in the K-K distance during this sister kinetochore movement in Δ SKAP kinetochore pairs (Figure S3). Notably, the K-K distance at the time of back kinetochore directional switching (starting to move away from the pole) was higher in Δ SKAP kinetochores than control (1.4±0.5 μm vs 1.0±0.2 μm, Figure 3F), again suggesting that Δ SKAP kinetochores are less sensitive to force changes. Together, these findings indicate that SKAP decreases friction between kinetochores and polymerizing microtubules.

**Figure 3.**
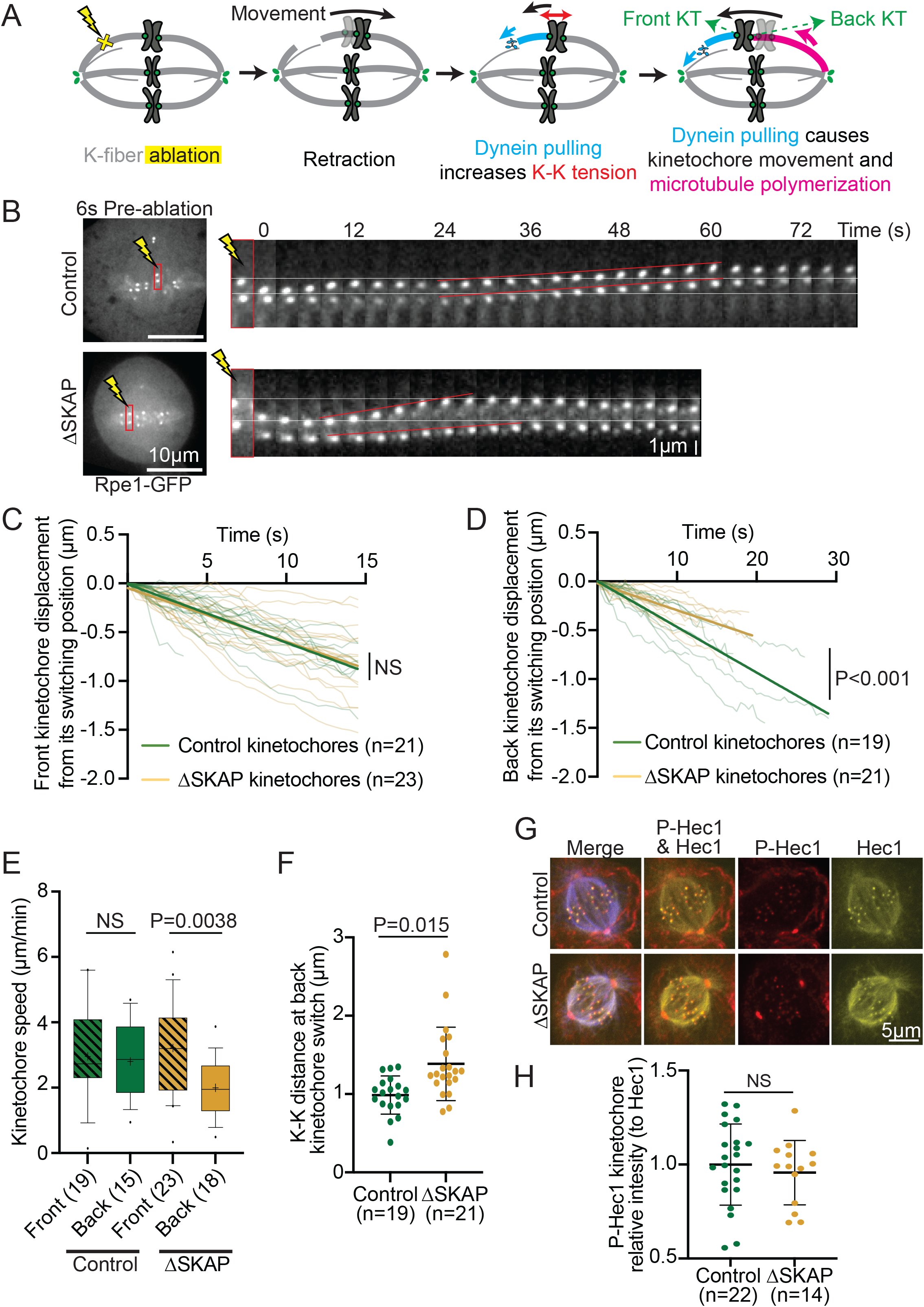
SKAP decreases kinetochore friction on polymerizing microtubules. (A) K-fiber ablation assay to isolate kinetochores associated to polymerizing microtubules: laser ablation (yellow X) leads to transient retraction of the kinetochore pair, the creation of new minus-ends leads to the recruitment of dynein (light blue) which pulls on a kinetochore and leads the microtubules attached to its sister to polymerize (pink). (B) Representative time lapse of k-fiber ablation in Rpe1-GFP control and ΔSKAP cells. White lines mark the kinetochore positions pre-ablation at the top, red lines represent the movement of sister kinetochores. (C) Individual tracks from control and ΔSKAP front kinetochores moving post-ablation, showing kinetochore displacement from its switching position after dynein pulling (time=0 corresponds to the timepoint of switch to poleward movement), with linear regression fits (straight lines) (analysis of covariance test, ANCOVA). (D) Individual tracks from control and ΔSKAP back kinetochores moving post-ablation, showing kinetochore displacement from its switching position after dynein pulling (time=0 corresponds to the timepoint of switch to away from the pole movement), with linear regression fits (straight lines) (analysis of covariance test, ANCOVA). (E) Front (lined boxes) and back (clear boxes) kinetochore speed distribution post-ablation in control and ΔSKAP kinetochores (student’
ss t-test) from data in (C) and (D). (F) K-K distance at the time the back kinetochore switched to away-from-pole movement post-ablation in control and ΔSKAP kinetochore pairs (Mann-Whitney test). In (C-F), n=number of ablations, one ablation per cell. (G) Representative immunofluorescence images of Rpe1-GFP control and ΔSKAP cells stained for Hec1-S69 phosphorylation (red), tubulin (blue) and Hec1 (yellow). (H) p-Hec1-S69 kinetochore intensity relative to Hec1 kinetochore intensity in control and ΔSKAP cells (student’s t-test; n=number of cells). See also Figure S3, Movies S3-4.

Hec1 dephosphorylation is known to increase friction (Zaytsev et al, 2014; Long et al, 2017), and Hec1-S69 phosphorylation maintains basal kinetochore movement dynamics at metaphase (DeLuca et al, 2018). Therefore, we tested whether SKAP depletion affected the level of Hec1-S69 phosphorylation. Immunofluorescence quantification showed that SKAP depletion did not affect kinetochore levels of Hec1-S69 phosphorylation (Figure 3G-H), indicating that friction regulation by SKAP occurs independently of Hec1-S69 phosphorylation. Together, these findings indicate that SKAP, in contrast to the proposed role of other microtubule binding proteins at the kinetochore (Zaytsev *et al*, 2014; Long *et al*, 2017; Helgeson *et al*, 2018; Huis In ’T Veld *et al*, 2019), decreases friction between kinetochores and polymerizing microtubules, and does so through a mechanism independent of that setting Hec1’s basal dynamic state (DeLuca et al, 2018). SKAP could either do so by increasing microtubule tip polymerization dynamics, producing an apparent change in friction, or by directly reducing kinetochore friction on the microtubule lattice.

### SKAP increases k-fiber depolymerization velocity and kinetochore force-responsiveness

Given that SKAP regulates the kinetochore’s frictional interface with polymerizing microtubules (Figure 3), we asked whether it also regulates its interface with depolymerizing ones. To decouple both attached sister kinetochores and isolate SKAP’s role on depolymerizing microtubules, we laser ablated one kinetochore in a sister pair in Rpe1-GFP cells, and tracked the movement of the remaining sister. As expected, the remaining sister was pulled poleward by depolymerizing microtubules, and reversed direction near the pole, pushed by polar ejection forces (McNeill & Berns, 1981; Khodjakov & Rieder, 1996; Ke *et al*, 2009) (Figure 4A-B, Movie S5). The poleward velocity of Δ SKAP kinetochores was slower than that in control (2.1±0.6 μm/min vs 3.5±0.8 μm/min, Figure 4B-D, Movie S5-6). This indicates that SKAP increases the velocity of microtubule depolymerization at the kinetochore interface; here too, it could do so either by lowering friction on the microtubule lattice or by increasing depolymerization dynamics and thereby decreasing apparent friction. Consistent with SKAP acting directly at the interface, SKAP depletion did not detectably change kinetochore levels of key microtubule plus-end depolymerases involved in metaphase kinetochore movements, MCAK and Kif18A (Wordeman *et al*, 2007; Stumpff *et al*, 2008) (Figure S4). Further, in the same kinetochore ablation experiments Δ SKAP kinetochores switched direction (to away from pole movement) closer to the pole than control kinetochores as they got pushed by polar ejection forces (3.2±1.2 μm vs 4.0±1.8 μm (Figure 4E)). Thus, SKAP increases the kinetochore’s force sensitivity when it is bound to depolymerizing microtubules, favoring a switch to the polymerization state, as observed after k-fiber ablation (Figure 3F) and during metaphase oscillations (Figure 1G). Together, these findings indicate that Astrin-SKAP does not increase the kinetochore’s grip on microtubules as other microtubule-binding proteins are thought to do, but instead reduces grip, lowering friction and effectively lubricating the interface, making it more dynamic and force responsive.

**Figure 4.**
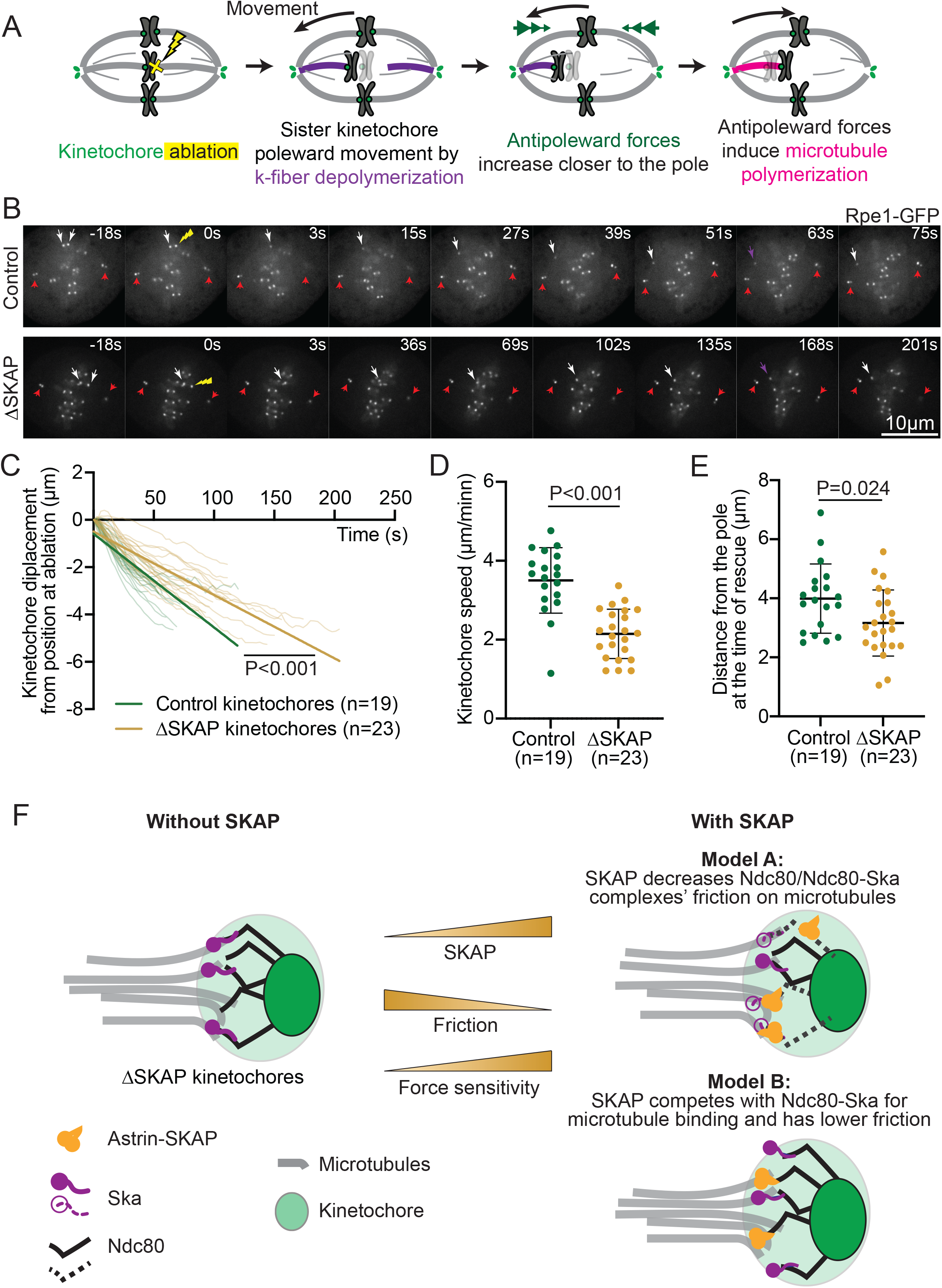
SKAP increases k-fiber depolymerization velocity and kinetochore force-responsiveness. (A) Kinetochore ablation assay to isolate kinetochores associated to depolymerizing microtubules: laser ablation (yellow X) of one sister leads to the other sister moving poleward as its microtubules depolymerize (purple), and to later move away-from-the pole as polar ejection forces (green arrowheads) increase and microtubules rescue and polymerize (pink). (B) Representative time lapse images of kinetochore ablation (yellow arrow) in control (top) and ΔSKAP (bottom) Rpe1-GFP cells, with remaining sister kinetochore (white arrow) and centrioles (red arrows) marked, and kinetochore directional switch marked (purple arrow). (C) Distance to position at ablation as a function of time for individual kinetochores post-ablation (t=0 corresponds to the first timepoint post-ablation) in control and ΔSKAP cells, with linear regression fits (straight lines) (analysis of covariance test, ANCOVA). (D) Average speed of individual kinetochores post sister ablation in control and ΔSKAP cells (Mann-Whitney test). (E) Kinetochore distance from the spindle pole of individual control and ΔSKAP cells at the time of direction switch from poleward to away-from-pole movement (rescue) post-ablation (student’
ss t-test), with a smaller distance typically reflecting a higher force at rescue. In (C-E), n=number of ablations, one ablation per cell. (F) Representation of the kinetochore-microtubule interface in the absence (left) and presence (right) of SKAP. Two models for how Astrin-SKAP (yellow) could increase friction at the kinetochore-microtubule interface (right). In Model A (top right), SKAP affects how Ndc80 (black) or Ndc80-Ska (purple) complexes bind microtubules (dashed black and purple), decreasing their friction on microtubules and indirectly reducing attachment friction and increasing dynamics. In Model B (bottom right), SKAP directly binds microtubules, with Ndc80-SKAP and Ndc80-Ska competing for microtubule binding with similar affinities (binding energy) but with Ndc80-SKAP moving on microtubules with lower friction (lower transition state energy in moving between lattice binding sites). In both models, the more SKAP molecules are at the kinetochore, the lower the friction at the kinetochore-microtubule interface and the higher the sensitivity to force (center). See also Figure S4, Movies S5-6.

## DISCUSSION

Faithful chromosome segregation requires kinetochores to hold on to dynamic microtubules that generate force. Here we ask: How does the mammalian kinetochore-microtubule interface stay dynamic and force responsive while maintaining a strong grip on microtubules? Mechanisms increasing grip as attachments mature are being actively studied, including Ndc80 dephosphorylation (Zaytsev *et al*, 2014; Long *et al*, 2017) and Ska recruitment (Auckland *et al*, 2017; Huis In ’T Veld *et al*, 2019). Here, we combine molecular and mechanical perturbations to define the mechanical role of SKAP at this interface, and we show that it in turn decreases grip at the interface. We demonstrate that SKAP increases sister kinetochore mobility and coordination in metaphase (Figure 1), and decreases tension at the kinetochore-microtubule interface (Figure 2). We show that SKAP increases the velocity at which kinetochores move on both polymerizing (Figure 3) and depolymerizing (Figure 4) microtubules, and that it makes the attachment more responsive to force changes (Figures 3-4). Together, our findings indicate that SKAP reduces friction at the kinetochore-microtubule interface, effectively lubricating it. As such, and given SKAP’s arrival as kinetochores biorient (Fang *et al*, 2009; Schmidt *et al*, 2010), we propose that SKAP keeps correct kinetochore-microtubule attachments dynamic to preserve them as they stabilize and mature (Zaytsev *et al*, 2014; Auckland *et al*, 2017). The association of SKAP mutations and changes in expression with some cancers and aneuploidy (Siprashvili *et al*, 2014; Jaju *et al*, 2015; Deng *et al*, 2021) is consistent with SKAP’s lubrication function being key for faithful chromosome segregation.

Our work indicates that SKAP lowers friction at the kinetochore-microtubule interface, raising the question of what function a lower friction could serve. Basal levels of attachment dynamics are essential for accurate chromosome segregation (Bakhoum *et al*, 2009; DeLuca *et al*, 2018). Ska recruitment and gradual dephosphorylation of Hec1 increases kinetochore grip on microtubules during mitosis (Zaytsev *et al*, 2014; Long *et al*, 2017; Auckland *et al*, 2017; Huis In ’T Veld *et al*, 2019). In contrast to other Hec1 phosphosites, S69 is persistently phosphorylated during mitosis, maintaining basal dynamics (DeLuca *et al*, 2018). Yet, these dynamics are lost and friction increases upon SKAP depletion (Figure 1, 3-4), although phosphorylation levels of Hec1-S69 are not affected by this depletion (Figure 3G-H). Thus, SKAP acts as a lubricant independently of Hec1-S69 phosphorylation. By reducing friction, SKAP may increase microtubule dynamics, increasing microtubule depolymerization (Figure 4B-D) and poleward flux (Wang *et al*, 2012). To our knowledge, SKAP is the first microtubule binder proposed to decrease friction at the kinetochore-microtubule interface. We propose that using SKAP to lower friction, instead of loosening Ndc80-Ska’s grip to lower friction, allows the cell to keep a dynamic kinetochore-microtubule interface without losing grip of microtubules and attachment stability, by having one molecule specialized for each activity. Indeed, having distinct mechanisms to tune grip and dynamics may lead to finer regulatory control “knobs” to preserve stable attachments.

By lubricating correct, bioriented attachments, SKAP could in principle help preserve them and prevent their increased microtubule affinity from making them less mobile and less responsive to force. The presence of more dynamic, lower friction interactions with microtubules could help stabilize attachments under force by increasing their adaptability to force changes. The force responsiveness of attachments is essential for accurate chromosome movement, alignment and sister kinetochore coordination (Ke *et al*, 2009; Stumpff *et al*, 2012; Wan *et al*, 2012), and SKAP depleted cells have chromosome alignment and segregation defects (Fang *et al*, 2009; Schmidt *et al*, 2010; Dunsch *et al*, 2011). Indeed, we show that SKAP makes kinetochores more mobile (Figure 1C-D), and the kinetochore-microtubule interface more sensitive to force changes (Figure 1E-G, 3F, 4E). This could explain why Δ SKAP attachments are under higher tension (Figure 2), inefficient at dissipating force (Figure 3B-E) and at responding to sister movement (Figure 1E-G, 3F), and it suggests that a function of lowering friction could be to make kinetochores responsive to force. While additional work will be needed to reveal how SKAP increases force responsiveness, in principle lowering friction would be sufficient to do so.

While SKAP reduces friction at the kinetochore-microtubule interface (Figure 3A-E), the mechanism by which it does so is not clear. SKAP depletion could in principle increase friction by reducing microtubule dynamics. However, lowering microtubule dynamics globally with drugs (as taxol or eribulin) decreases interkinetochore tension (or K-K distance), rather than increase it as we see for Δ SKAP (Figure 2) (Waters *et al*, 1998; Okouneva *et al*, 2008). Thus, decreasing microtubule dynamics is not sufficient to recapitulate the effects of SKAP depletion. Further, we could not detect any change in kinetochore recruitment of key microtubule dynamics regulators (MCAK, Kif18A) in Δ SKAP cells (Figure S4) and changing their activity is not consistent with our findings: MCAK depletion decreases both interkinetochore tension and microtubule depolymerization (Wordeman *et al*, 2007), and Kif18A depletion increases kinetochore velocity and oscillations amplitudes and decreases switching rates (Stumpff *et al*, 2008, 2012), and we do not see these observables change as such (Figures 1C-D, 2, 3F, 4C-E). While we cannot exclude the possibility that SKAP regulates the kinetochore activity of these or other microtubule dynamics regulators, a change in microtubule dynamics alone is not sufficient to recapitulate all the behaviors of Δ SKAP cells. Together, these findings are consistent with SKAP playing a mechanical role at the kinetochore-microtubule interface, and not simply regulating microtubule dynamics.

Mechanistically, SKAP could regulate kinetochore-microtubule friction in different ways. One way it could do so is by changing the sliding friction of other kinetochore proteins on microtubules. SKAP could induce conformational changes in Ndc80 or Ska complexes which lower their friction on microtubules (Model A, Figure 4F), independently of Hec1-S69 phoshoregulation (Figure 3G-H). Alternatively, SKAP could directly bind microtubules and act as a kinetochore-microtubule coupler, as suggested by in vitro work (Schmidt *et al*, 2010; Friese *et al*, 2016; Kern *et al*, 2017). As such, it could then compete out other microtubule binders that have higher friction on microtubules (Model B, Figure 4F), as microtubule affinity (binding energy) and friction coefficient (transition state energy in moving between lattice binding sites) are not strictly coupled (Forth *et al*, 2014). The affinity of Ndc80-SKAP on microtubules (Kern *et al*, 2017) is similar, though lower, than that of Ndc80-Ska (Schmidt *et al*, 2012), suggesting that both complexes could compete for microtubule binding; if Ndc80-SKAP had lower friction on microtubules than Ncd80-Ska, SKAP kinetochore localization could lower friction at the kinetochore-microtubule interface. In principle, the above models hold independently of how SKAP is recruited to the kinetochore, for example through Ndc80 (Schmidt *et al*, 2010; Kern *et al*, 2017) or Mis13 (Wang *et al*, 2012). Whether SKAP lowers other kinetochore proteins’ (Ndc80, Ska) friction on microtubules (Model A) and whether SKAP and other kinetochore proteins compete for microtubule binding and generate different friction (Model B) are non-exclusive models and a rich area of future study.

Across scales in biology, diverse interfaces need to be both strong and dynamic. For example, motor proteins must walk on their tracks without letting them go, and cell-cell junctions must robustly persist yet remodel. Understanding the mechanics of such interfaces will require not only understanding how strong, robust interactions are achieved, but how the interface can remain dynamic. The latter may require looking beyond mechanisms that grip, as is the case for SKAP which makes the human kinetochore-microtubule interface more dynamic.

## Supplementary Figure Legends

**Figure S1.**
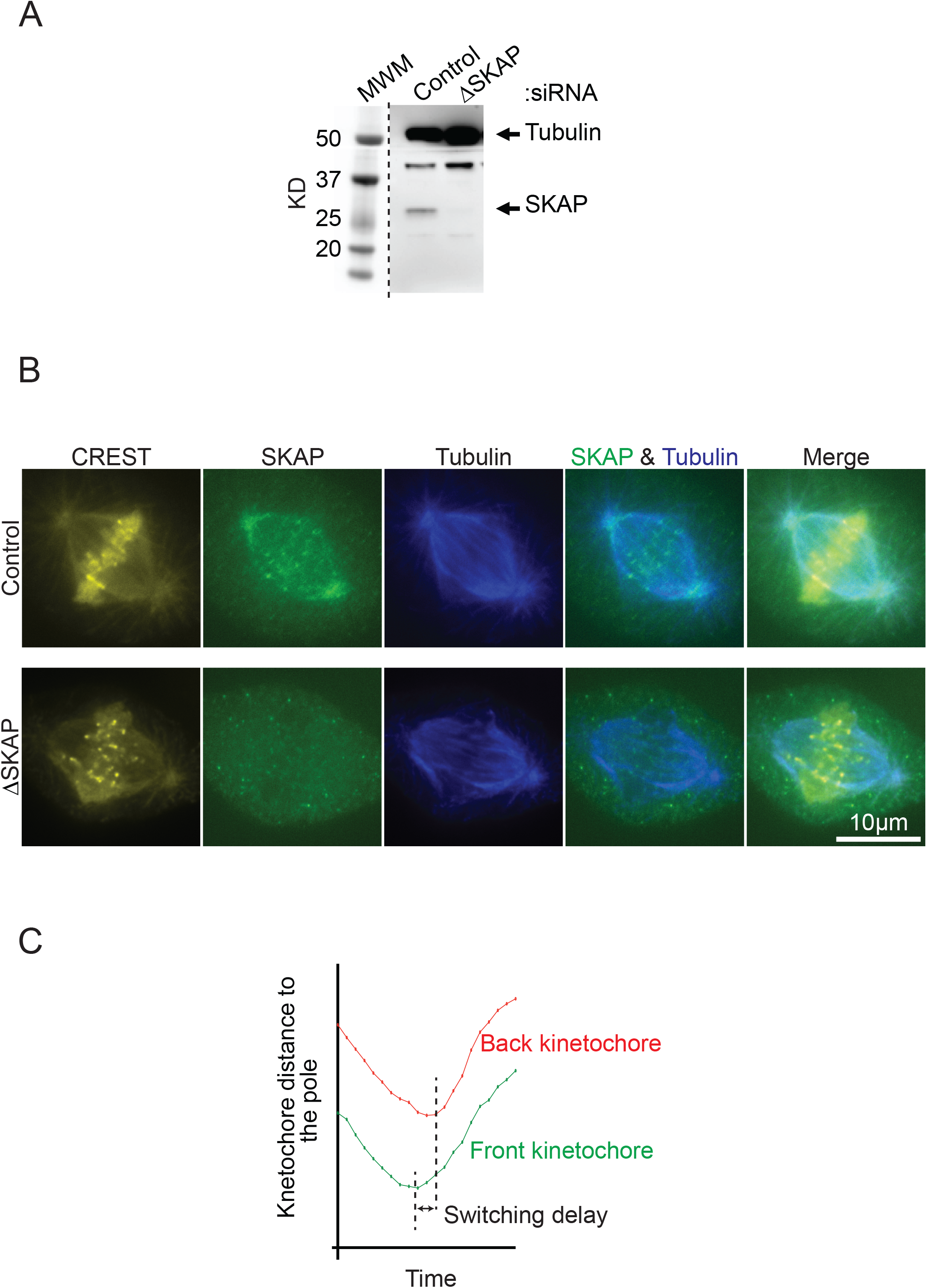
SKAP silencing and kinetochore oscillations. Related to Figure 1. (A) Representative western blot of SKAP expression in control and ΔSKAP in Rpe1-GFP cells at 24 h post-transfection (80-95% silencing), with tubulin as a loading control and with molecular weight marker (MWM). (B) Representative immunofluorescence images in control and ΔSKAP Rpe1-GFP cells (24 h post-transfection) stained for CREST (yellow), tubulin (blue) and SKAP (green). (C) Schematic representation of how sister kinetochores typically switch direction during metaphase oscillations (Wan *et al*, 2012).

**Figure S2.**
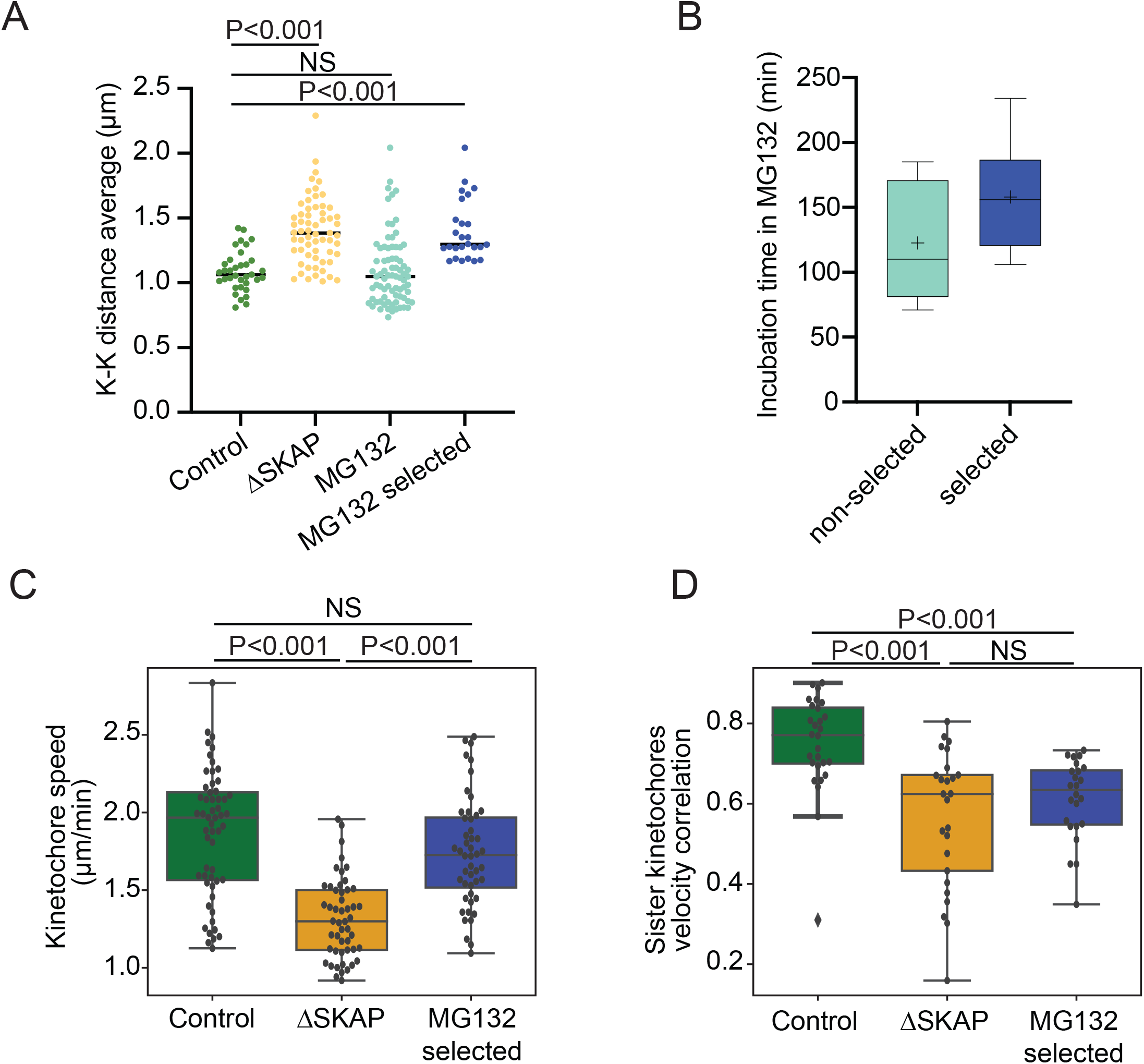
SKAP depletion is not consistent with a decrease in centromere stiffness. Related to Figure 2. (A) Average K-K distance for individual sister kinetochore pairs over time in live control (n=34 pairs from 18 cells), ΔSKAP (n=61 from 20 cells), MG132-treated (n=77 pairs from 18 cells), and MG132-treated “selected” (n=29 pairs from 6 cells) Rpe1-GFP cells (1-4 pairs per analyzed cell). The latter “selected” pairs only include those with mean K-K distances higher than two standard deviations over average control pairs. (B) Time of MG132 incubation of selected vs non-selected cells in (A). (C) Individual kinetochore speed during metaphase oscillations in control, ΔSKAP or MG132-treated selected cells (Mann-Whitney test) for the cells from (A). (D) Sister kinetochore velocity correlation during metaphase oscillations in control, ΔSKAP and MG132-treated selected cells from (A) (Mann-Whitney test).

**Figure S3.**
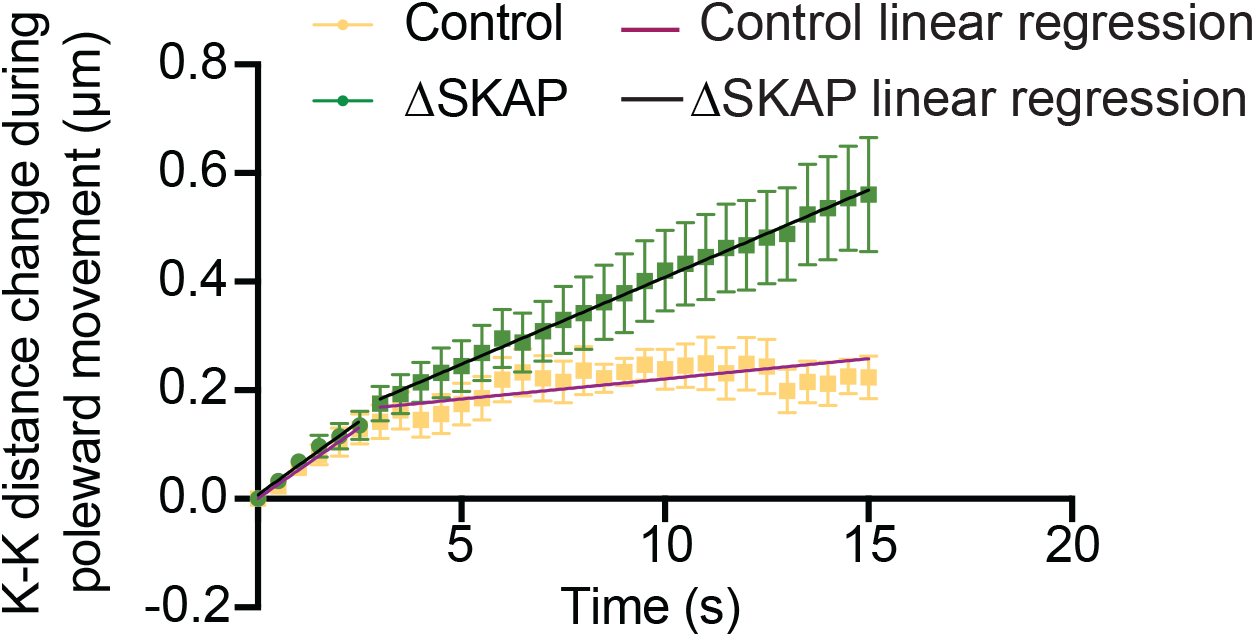
Differential velocity between front and back ΔSKAP sister kinetochores leads to increasing K-K distance after k-fiber ablation. Related to Figure 3. K-K distance during front kinetochore poleward movement after k-fiber ablation in Rpe1-GFP control and ΔSKAP cells, with t=0 at the start of front kinetochore poleward movement. Linear regression slope is similar in control and ΔSKAP cells during the first 2.5 s, but not at later timepoints (analysis of covariance test, ANCOVA). Tracks from Figure 2B dataset.

**Figure S4.**
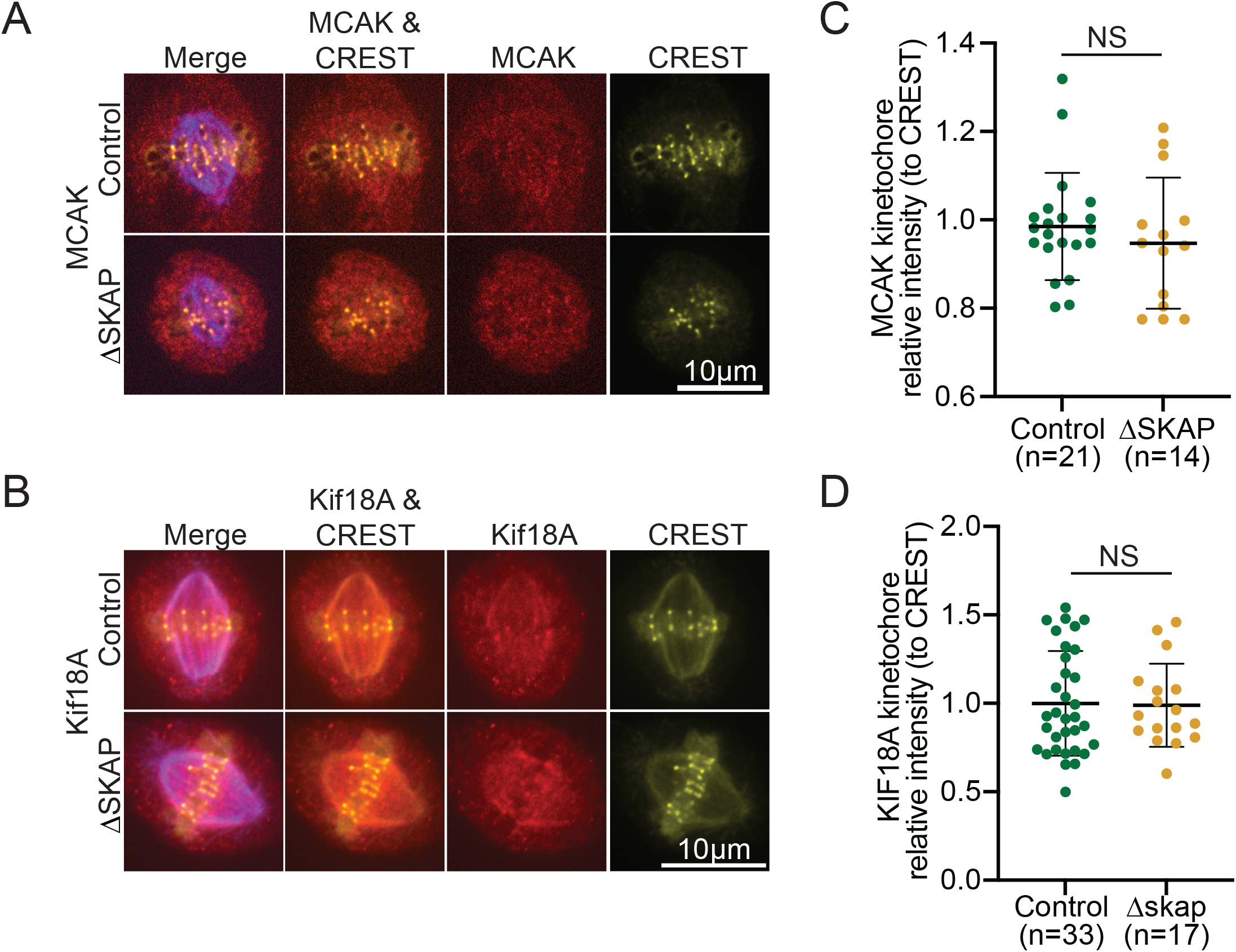
SKAP does not affect kinetochore recruitment of two proteins (MCAK and Kif18A) regulating microtubule dynamics. Related to Figure 4. Representative immunofluorescence images in control and ΔSKAP Rpe1-GFP metaphase cells with staining for tubulin (purple), CREST (yellow), and A) MCAK (red) or B) Kif18A (red). Kinetochore intensity of (C) MCAK and (D) Kif18A in control and ΔSKAP cells relative to CREST kinetochore intensity (student’
ss t-test; n=number of cells), obtained from immunofluorescence images as in (A) and (B), respectively.

## Movie Legends

**Movie S1. Metaphase kinetochore oscillations in control cell**. Related to Figures 1-2, representative kinetochore oscillations in a metaphase control Rpe1-GFP cell. Kinetochores (GFP-CenpA) and centrioles (centrin1-GFP) in grey. Time in min:s. The movie was collected using a spinning disk confocal microscope at 1 frame every 3 s and adjusted to play at 75x real time. Movie corresponds to cell shown in Figure 1B.

**Movie S2. Metaphase kinetochore oscillations in Δ SKAP cell**. Related to Figures 1-2, representative kinetochore oscillations in a metaphase Δ SKAP Rpe1-GFP cell. Kinetochores (GFP-CenpA) and centrioles (centrin1-GFP) in grey. Time in min:s. The movie was collected using a spinning disk confocal microscope at 1 frame every 3s and adjusted to play at 75x real time. Movie corresponds to cell shown in Figure 1B.

**Movie S3. K-Fiber ablation in control cell**. Related to Figure 3. Laser ablation of a k-fiber in a metaphase control Rpe1-GFP cell. Kinetochores (GFP-CenpA) and Centrioles (centrin1-GFP) in grey, time in seconds. After k-fiber ablation (yellow arrow, time = 0) and immediate K-K distance relaxation, the front kinetochore (sister kinetochore on the ablation side) is pulled poleward by dynein activity at the new k-fiber minus-ends. K-K distance increases as a result of this force, triggering microtubule polymerization at the back k-fiber and back kinetochore (away-from-pole) movement (following the front one). The movie was collected using a spinning disk confocal microscope at 1 frame every 3 s before ablation and 0.5 s after ablation and adjusted to play at 8 frames/s. Movie corresponds to cell shown in Figure 3B.

**Movie S4. K-fiber ablation in Δ SKAP cell**. Related to figure 3. Laser ablation of a k-fiber in a metaphase Δ SKAP Rpe1-GFP cell. Kinetochores (GFP-CenpA) and centrioles (centrin1-GFP) in grey, time in seconds. After k-fiber ablation (yellow arrow, time = 0) and immediate K-K distance relaxation, the front kinetochore (sister kinetochore on the ablation side) is pulled poleward by dynein activity at the new k-fiber minus-ends. K-K distance increases as a result of this force, triggering microtubule polymerization at the back k-Fiber and back kinetochore away-from-pole movement (following the front one). The movie was collected using a spinning disk confocal microscope at 1 frame every 3 s before ablation and 0,5 s after ablation and adjusted to play at 8 frames/s. Movie corresponds to cell shown in Figure 3B.

**Movie S5. Kinetochore ablation in control cell**. Related to Figure 4. Laser ablation of a kinetochore in a metaphase control Rpe1-GFP cell. Kinetochores (GFP-CenpA) and Centrioles (centrin1-GFP) in grey, time in seconds. After individual kinetochore ablation (yellow arrow, time = 0), the sister kinetochore moves poleward, driven by microtubule depolymerization at the associated k-fiber. Kinetochore movement switches to away-from-pole direction (purple arrow) near the spindle pole due to polar ejection forces that trigger microtubule rescue. The movie was collected using a spinning disk confocal microscope at 1 frame every 6 s before ablation and 3 s after ablation and adjusted to play at 10 frames/s. Movie corresponds to cell shown in Figure 4B.

**Movie S6. Kinetochore ablation in Δ SKAP cell**. Related to Figure 4. Laser ablation of a kinetochore in a metaphase Δ SKAP Rpe1-GFP cell. Kinetochores (GFP-CenpA) and Centrioles (centrin1-GFP) in grey, time in seconds. After individual kinetochore ablation (yellow arrow, time = 0), the sister kinetochore moves poleward, driven by microtubule depolymerization at the associated k-fiber. Kinetochore movement switches to away-from-pole direction (purple arrow) near the spindle pole, due to polar ejection forces that trigger microtubule rescue. The movie was collected using a spinning disk confocal microscope at 1 frame every 6 s before ablation and 3 s after ablation and adjusted to play at 10 frames/s. Movie corresponds to cell shown in Figure 4B.

## Methods

### Cell culture, siRNA transfection and drug treatments in Rpe1 cells

Rpe1-GFP cells (gift from A. Khodjakov, Wadsworth Center (Paul *et al*, 2011)) were cultured in DMEM/F12 (Thermo Fisher Scientific, 11320082) supplemented with 10% qualified and heat-inactivated fetal bovine serum (FBS) (10438-026, Gibco) and penicillin/streptomycin and maintained at 37°C and 5% CO_2_. Cells were plated in 35 mm glass-bottom dishes (poly-D-lysine coated; MatTek Corporation) for live imaging experiments or in six wells plates after addition on #1.5 25 mm coverslips (acid cleaned and poly-L-lysine coated) for immunofluorescence experiments. For knockdown experiments, siRNA targeting SKAP (5′-AGGCUACAAACCACUGAGUAA-3′) or luciferase siRNA (5’ CGUACGCGGAAUACUUCGA 3’, control) were transfected in Rpe1-GFP cells and, using 4 µl of lipofectamine siRNAmax (Thermo Fisher Scientific, 13778075) and 10 µM of siRNA for 2 ml of cell culture media. Cells were incubated at 37 °C for 6-8 h before media wash, and imaged or processed 24 h after transfection. SiR-tubulin (Cytoskeleton, CY-SC002) was used at 1 µM concentration and added to the cells 40 min before imaging in experiments for Figure 2C. Nocodazole (Sigma-Aldrich, M1404-50MG) was used at 2 µM concentration and added 3 h before fixation in experiments for Figure 2 D-E. MG132 (EMB Millipore, 474790-5MG) was used at 10uM concentration in experiments for Figure S2.

### Immunofluorescence and immunoblotting

To validate the SKAP siRNA (Figure S1A-B), cells were seeded in six well plates, transfected with luciferase (control) or SKAP siRNA 24 h after transfection and processed for immunoblotting. Cells were collected in PBS1X using a cell scrapper and lysed in PBS1X + 1% NP40 on ice for 30 min. Samples were run in a 4-12% Bis-Tris gel (Invitrogen, NPO335BOX) and transferred to a nitrocellulose membrane (Thermo Scientific, 88018). The following primary and secondary antibodies and dyes where used (incubated in TBS1X, 3 % milk, 0.1 % Tween for 1 h or 45 min, respectively): anti-SKAP (1ug/ml, rabbit, Origene, TA333584), anti-α-tubulin (DM1A, 1:1000, mouse, Sigma, T6199), goat anti-mouse IgG-HRP (1:1000, sc-2005, Santa Cruz Biotechnology) and mouse anti-rabbit IgG-HRP (1:1000, sc-2357, Santa Cruz Biotechnology). Blots were exposed with SuperSignal West Pico Substrate (Thermo Scientific) and imaged with a Bio-Rad ChemiDoc XRS+ system.

For immunofluorescence experiments cells were fixed in 99.8% methanol for 10 min at -20 °C, 24 h after siRNA transfection, and permeabilized in PBS1x, 0.5% BSA, 0.1% Triton (IF buffer thereafter) for 30 min. The following primary antibodies were incubated for 1 h in IF buffer: SKAP (1ug/ml, rabbit, gift from I. Cheeseman (Kern *et al*, 2016)), Kif18A (2ug/ml, rabbit, Bethyl, A301-080A-M), MCAK (1:500, rabbit, cytoskeleton, AKIN05), α-Tubulin DM1A (1:1000, mouse, Sigma, T6199), p-Hek1-s69 (1:3000, rabbit, gift from J. DeLuca (DeLuca *et al*, 2018)), CREST (1:100, human, Antibodies Incorporated, 15-234-0001), Hec1 (1:200, mouse, Abcam, ab3613), tubulin (1:2000, rat, Bio-rad, MCA77G). There washes in IF buffer (10 min each) were done before incubation with the following secondary antibodies (1:1000 in IF buffer, 45 min): goat anti-mouse IgG Alexa Fluor 405, 488 and 568 (A31553, A11001 and A11004, Invitrogen), goat anti-rabbit IgG Alexa Fluor 405 and 568 (A31556 and A11011, Invitrogen), goat anti-rat IgG Alexa Fluor 488 (A11006, Invitrogen) and goat anti-human IgG Alexa Fluor 568 (A21090, Invitrogen). Samples were washed once in IF buffer and twice in PBS1X (10 min each) before mounting in ProLong Gold Antifade reagent (P36934, Thermo Fisher).

### Microscopy and laser ablation

Samples were imaged using an inverted microscope (Eclipse Ti-E; Nikon) with a spinning disk confocal (CSU-X1; Yokogawa Electric Corporation), head dichroic Semrock Di01-T405/488/568/647 for multicolor imaging, equipped with 405 nm (100 mW), 488 nm (120mW), 561 nm (150mW), and 642 nm (100mW) diode lasers, emission filters ET455/50M, ET525/ 50M, ET630/75M and ET690/50M for multicolor imaging, and an iXon3 camera (Andor Technology) operated by MetaMorph (7.7.8.0; Molecular Devices) (Elting *et al*, 2014). Cells were imaged through a 100X 1.45 Ph3 oil objective and 1.5X lens.

For live imaging and laser ablation experiments, cells were maintained in a stage-top incubation chamber (Tokai Hit) at 37 °C and 5 % CO_2_. Metaphase oscillations were imaged every 3 s (Figures 1, 2B and S2). Laser ablation (30-40 pulses of 3 ns at 20 Hz) with 514 nm light was performed using the MicroPoint Laser System (Andor). For laser ablation experiments, images were acquired more slowly prior to ablation and then acquired more rapidly after ablation (3 s prior and 0.5 s after k-fiber ablation, and 6 s prior and 3 s after kinetochore ablation (Figures 3 and 4, respectively)). Successful k-fiber ablation was verified by immediate K-K distance relaxation (Figure 2G-I, 3 and S3) and posterior front kinetochore movement poleward by dynein pulling. Successful kinetochore ablation was verified by immediate poleward movement of the remaining sister kinetochore (Figure 4). For long term imaging experiments (Figure 2C), Rpe1-GFP cells treated with 100 nM SiR-DNA were imaged every 4 min for 18-20 h using a 20X objective.

### Study design and data inclusion criteria

Two general criteria for inclusion of cells in metaphase oscillation and laser ablation experiments were applied. First, cells must express detectable levels of GFP-CenpA at kinetochores, but not so high as to completely label chromosome arms. Second, cells must be in metaphase, with a defined metaphase plate. For oscillation experiments (Figure 1-2B and S2), 2-4 kinetochore pairs per cell were analyzed, and the two kinetochores from the pair must stay in focus for a minimum of 90 s. For MG132 treatment experiments (Figure S2), cells with low centromere stiffness where selected if their mean K-K distance over time was higher than two standard deviations over the mean control K-K distance. For kinetochore speed calculations in ablation experiments (Figure 3E-4D), kinetochore tracks shorter than 5 timepoints were excluded due to the absence of a consistent directional movement.

We did not pre-estimate a required sample size before performing experiments nor did we blind or randomize samples during experimentation or analysis. The ablation experiments in this study are low throughput by nature, which does not enable us to report averages from multiple independent replicate experiments. Instead, we pool cells from across different independent experiments (with at least three independent experiments per condition per assay).

### Analyzing fluorescence intensity

Images were processed and fluorescence intensity was quantified using FIJI (Version 2.3.0/1.53f) (Schindelin *et al*, 2012). For protein intensity measurements, a color threshold mask (Yen method) was applied using the CREST or Hec1 signal (for kinetochore selection) or DM1A signal (for spindle microtubule selection) to define the areas in which the fluorescence intensity would be measured for each protein of interest, and this intensity was normalized by the kinetochore or microtubule marker intensity. Figures 3H and S4C-D showing data from individual representative experiments, three independent experiments were performed for each quantification, obtaining comparable results.

### Analyzing kinetochore behaviors

Kinetochores and centrioles were manually tracked from GFP-CenpA /centrin1-GFP movies using the MtrackJ plugin from FIJI. GFP-centriole position was used as a marker for the spindle pole position. Kinetochore position was calculated as the distance from the spindle pole position. In metaphase oscillations experiments, all quantifications and statistical analyses were performed using home-written Phyton code. Kinetochore speed at each timepoint was calculated as the difference in kinetochore position between two consecutive timepoints. Kinetochore speed was calculated by obtaining the slope of the best fitting regression line of individual kinetochore tracks (Figures 3E and 4D) or of entire tracks together (Figures 3C, D and 4C). Sister kinetochore movement coordination was obtained by calculating the correlation of sister kinetochore velocity over time or by the percentage of timepoints in which sister kinetochore movement direction was opposite. Kinetochore directional switch was determined by the consistent movement of a kinetochore in a the new direction for 3 consecutive timepoints. In ablation experiments, time after ablation was measured from the fist timepoint immediately after ablation. In all experiments, a kinetochore directional switch was defined by the consistent movement of the kinetochore in the opposite direction for a period of 3 consecutive timepoints (9 s in oscillations and kinetochore ablations and 1.5 s for k-fiber ablation experiments). K-K distance was calculated by subtracting the position of sister kinetochore A from sister kinetochore B, obtaining the length of the vector, and calculating the K-K distance average over time for each kinetochore pair. K-K distance relaxation was obtained by subtracting the K-K distance at the first timepoint after ablation from the last timepoint before ablation.

### Movie preparation

Movies were prepared using FIJI. Brightness and contrast were linearly adjusted to clearly visualize the kinetochores and centrioles.

### Statistical analysis

Statistical analysis was performed in Phyton or Graphpad (Prism 9). The Fisher’s exact test was used in Figure 1G, Student’s T test was used for parametric datasets (Figures 2B,I, 3E,H, 4E ad S4C-D), Mann-Whitney test for non-parametric datasets (Figures 1C-F, 2C,E, 3F, 4D and S2A,C-D) and analysis of covariance test, ANCOVA, for linear regression slopes comparison (Figures 3C,D-4C and S3).

## Author contributions

Conceptualization, M.R.S., S.D.; Methodology, M.R.S.; Software, M.R.S., P.S.; Validation, M.R.S.; Formal analysis, M.R.S.; Investigation, M.R.S., R.S. (Figures 2C, 3G, S4); Resources, S.D.; Data curation, M.R.S.; Writing-original draft, M.R.S.; Writing-Review & Editing, M.R.S., S.D.; Visualization, M.R.S.; Supervision, M.R.S., S.D.; Funding Acquisition, S.D.

## Acknowledgments

We thank Iain Cheeseman for anti-SKAP antibody, Jennifer DeLuca for anti-Hec1-pS69 antibody, Alexey Khodjakov for Rpe1-GFP (GFP-CenpA and centrin1-GFP) cells and Rob Phillips for discussions. We thank the members of the Dumont Lab for discussions and critical reading of the manuscript. This work was supported by NIH DP2GM119177, NIH R01GM134132, NIH R35GM136420, NSF CAREER 1554139, and NSF 1548297 Center for Cellular Construction, the Rita Allen Foundation and the Chan Zuckerberg Biohub (S.D.).

